# A Biparatopic Intrabody Renders Vero Cells Impervious to Ricin Intoxication

**DOI:** 10.1101/2024.07.02.601761

**Authors:** Timothy F. Czajka, David J. Vance, Renji Song, Nicholas J. Mantis

**Affiliations:** Department of Biomedical Sciences, University at Albany, Albany, NY 12201 United States; Division of Infectious Diseases, Wadsworth Center, New York State Department of Health, Albany, NY, 12208; Division of Research, Wadsworth Center, New York State Department of Health, Albany, NY, 12208

**Keywords:** toxin, ribosome, single domain antibody, toxin, intrabody, antibody, neutralization, intracellular, biodefense

## Abstract

Expression of camelid-derived, single-domain antibodies (V_H_Hs) within the cytoplasm of mammalian cells as “intrabodies” has opened-up novel avenues for medical countermeasures against fast-acting biothreat agents. In this report, we describe a heterodimeric intrabody that renders Vero cells virtually impervious to ricin toxin (RT), a potent Category B ribosome-inactivating protein (RIP). The intrabody consists of two structurally defined V_H_Hs that target distinct epitopes on RT’s enzymatic subunit (RTA): V9E1 targets RTA’s P-stalk recruitment site, and V2A11 targets RTA’s active site. Resistance to RT conferred by the biparatopic V_H_H construct far exceeded that of either of the V_H_Hs alone and effectively inhibited all measurable RT-induced cytotoxicty *in vitro*. We propose that targeted delivery of bispecific intrabodies to lung tissues may represent a novel means to shield the airways from the effects of inhalational RT exposure.

## Introduction

The development of medical countermeasures (MCM) against the panel of biothreat agents and toxins, as defined by the Centers for Disease Control and Prevention more than 20 years ago, remains far from complete (Cieslak et al., 2018). One such case is ricin toxin (RT), a potent plant-derived, ribosome-inactivating protein (RIP) that provokes pulmonary congestion, edema, neutrophil infiltration, and, potentially, acute respiratory distress upon inhalation (Gal et al., 2017; Pincus et al., 2015; Roy et al., 2019; Roy et al., 2020). RT consists of two subunits, RTA and RTB, each with essential roles in promoting cell death (Montfort et al., 1987; Olsnes et al., 1974). RTB is a Gal/GalNAc-specific lectin that mediates uptake and retrograde trafficking of RTA from the plasma membrane to the endoplasmic reticulum (ER) (Sowa-Rogozinska et al., 2019). RTA is an RNA N-glycosidase that when translocated across the ER membrane into the cytoplasm catalyzes the hydrolysis of a single base within the sarcin-ricin loop (SRL) of the 28S rRNA, thereby arresting protein synthesis and triggering the so-called ribotoxic stress response (RSR) (Endo and Tsurugi, 1987; 1988; Snieckute et al., 2022). While we have described monoclonal antibodies (MAbs) capable of neutralizing RT and protecting non-human primates against RT intoxication when given prophylactically or within hours after exposure (Roy *et al*., 2019; Roy *et al*., 2020), there are currently no antidotes to neutralize RTA once it has gained access to the cell cytoplasm.

Camelid-derived, single-domain antibodies (sdAbs) have emerged as potent and highly specific neutralizing agents for a range of plant and bacterial toxins. sdAbs generally constitute the variable only region a class of camelid-specific, heavy chain-only antibodies (V_H_Hs) with unique intrinsic properties that have led to their widespread applications across the biomedical sciences (Muyldermans, 2021). V_H_Hs are small (∼15 kDa), conformationally stable immunoglobulin domains amenable to both expression in *Escherichia coli* and display on the surface of M13 phage (“phage display”). V_H_H phage display combined with reiterative rounds of screening (panning) against antigens of interest can yield diverse collections of antibodies with a range of epitope specificities, as evidenced by the ∼800 sdAbs directed against SARS-CoV-2 currently in the Coronavirus Antibody Database (Raybould et al., 2021). Moreover, V_H_HS can be linked together as homodimers, heterodimers or higher order multimers resulting in antibodies with increased avidity and breadth of activity (Laursen et al., 2018).

Over the course of the past ten years, dozens of RT-specific alpaca- and llama-derived V_H_Hs have been described in the literature (Anderson et al., 2008; Czajka et al., 2022; Herrera et al., 2016; Poon et al., 2017; Rudolph et al., 2020; Vance et al., 2020; Vance et al., 2013). In fact, at this point, RT-specific V_H_Hs have been identified that are capable of interfering with virtually every step in RT’s cytotoxic pathway, including cell attachment (Rudolph et al., 2021; Vance *et al*., 2020), retrograde trafficking (Rudolph *et al*., 2021), ribosome recruitment (Czajka and Mantis, 2022), and SRL depurination (Rudolph *et al*., 2020). Moreover, a subset of V_H_Hs were identified to be capable of neutralizing RT intracellularly when expressed as intrabodies (Czajka and Mantis, 2022; Rudolph *et al*., 2020). Of particular interest are two structurally well-characterized V_H_Hs, V2A11 and V9E1, each of which play a role in shielding the ribosome from RTA. V2A11 targets RTA’s active site, a solvent-exposed cleft on one face of the molecule that accommodates the protruding adenine (A) within the conserved GAGA motif of the mammalian SRL (Mlsna et al., 1993; Rutenber et al., 1991). Structural analysis revealed that V2A11’s complementarity-determining region 3 (CDR3) penetrates the active site pocket and interacts with catalytic residues Tyr80 and Tyr123, thereby obstructing any possible interaction with the SRL (Rudolph *et al*., 2020). By contrast, V9E1 engages with RTA’s ribosomal recruitment site (RRS), a hydrophobic pocket near the C-terminus of RTA that is normally masked by RTB (Chiou et al., 2008; Czajka and Mantis, 2022; Li et al., 2009; May et al., 2012). When RTA is translocated into the cytoplasm, the RRS is exposed and able to engage with the ribosomal P-stalk proteins (Li et al., 2018). Structural analysis of V9E1 reveals CDR3 occupancy of the RRS and antibody mimicry of the P-stalk protein interactions (Czajka and Mantis, 2022). When transiently expressed as intrabodies in the cell cytoplasm of Vero cells, V9E1 and V2A11 each conferred a high degree of resistance to RT-induced cytotoxicity (Czajka and Mantis, 2022; Rudolph *et al*., 2020). In this report, we sought to investigate whether a biparatopic antibody targeting of RTA’s active site (AS) with V2A11 and RTA’s P-stalk ribosome recruitment site (RRS) with V9E1 would confer a degree of protection against RT not achievable with either of the antibodies independently.

## Results

To evaluate the neutralizing activity of V2A11 and V9E1 alone and in combination, Vero cells were transfected with plasmids encoding V2A11, V9E1 or an equal mixture of the two, and then challenged one day later with escalating amounts of RT. In parallel studies we confirmed that Vero cell transfection with the indicated plasmids resulted in detectable levels of V2A11 and V9E1 in cell lysates (**Figure S1**). The amount of RT (ng/mL) required to reduce Vero cell viability by 50% at 48 h was considered the **EC**_**50**_. The resistance index (**RI**) was defined as the EC_50_ of V_H_H-transfected cells divided by EC_50_ of controls.

Cells transfected with V2A11 and V9E1 individually had EC_50_ values of 140 ng/mL and 700 ng/mL, respectively, as compared to 4.8 ng/mL for controls (**Figure 1**). This translates to RI values of ∼29 for V2A11 and ∼146 for V9E1. Vero cells dually transfected with V2A11- and V9E1-encoding plasmids had an EC_50_ value of approximately 7 ug/mL, an RI value of 1460, 10-fold or more resistant to RT as compared to either V_H_H alone, suggesting synergy between V9E1 targeting the RTA’s P-stalk binding site and V2A11 targeting RTA’s active site (**Figure 1**). To discern whether the hyper-resistance to RT was epitope specific or just a consequence of targeting two distinct regions on RTA, we doubly transfected Vero cells with V2A11 and V9F6, a V_H_H that recognizes an epitope immediately adjacent to (but not occluding) RTA’s P-stalk binding site (Czajka *et al*., 2022). Dual transfection of Vero cells with V2A11 and V9F6 resulted in EC_50_ and RI values that were less than V2A11 alone, indicating that co-transfection of V9F6 does not augment an V2A11’s neutralizing activity (**Figure S2**). Thus, the enhanced intracellular neutralization of RT by the combination of V2A11 and V9E1 is specific and not simply the result of two different intrabodies.

**Figure 1.**
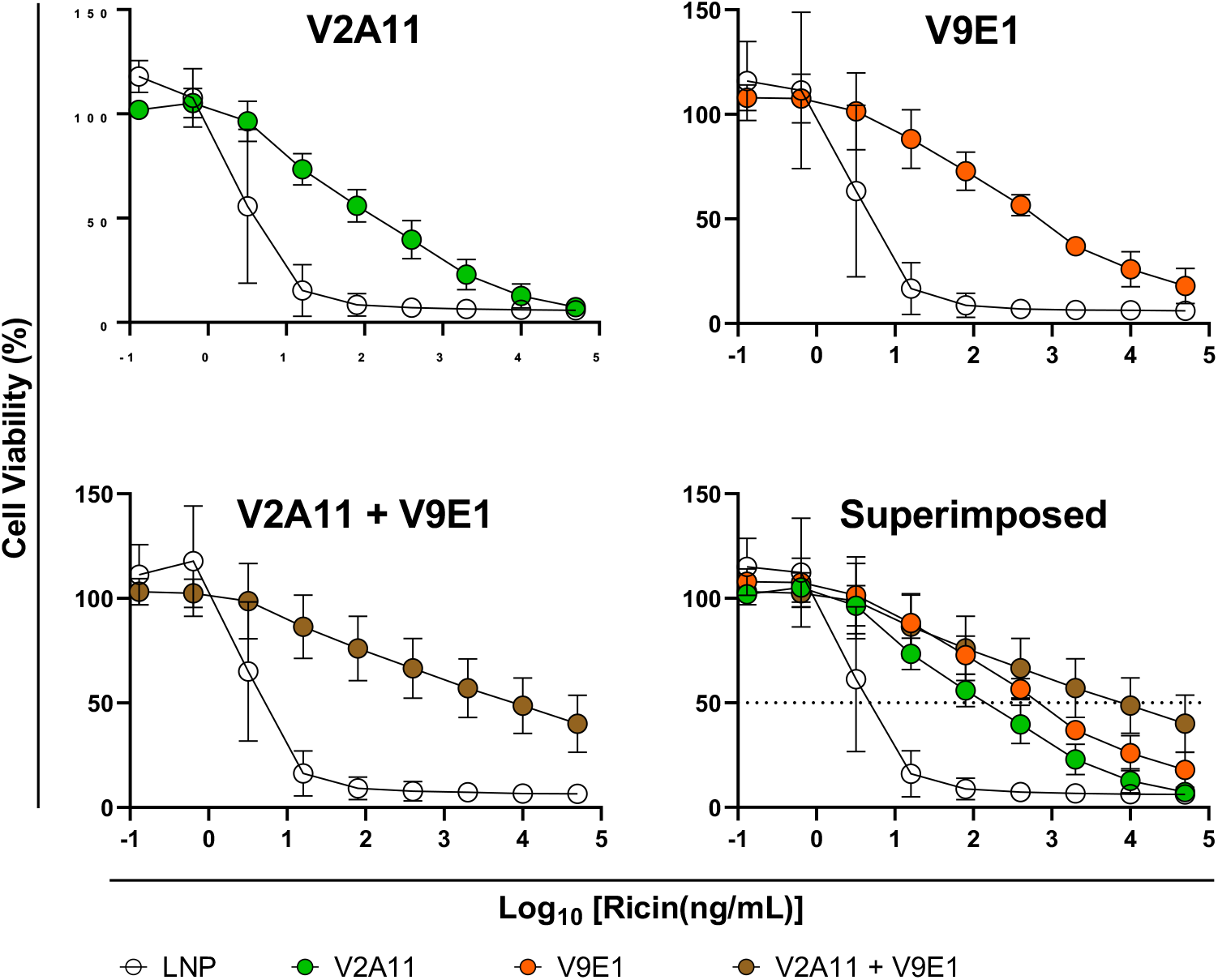
Intracellular neutralization of ricin by co-expressed neutralizing intrabodies. V2A11 (green; top left), V9E1 (orange; top right), or a combination of both intrabodies (brown; bottom left) were transfected into Vero cells and treated with a range of ricin concentrations one day later. A superposition of all three is shown bottom right. Cell viabilities were analyzed as a percentage of control live cells (not treated with ricin) two days after intoxication, and compared to cell transfected with vehicle LNP controls (white).

We next postulated that physically linking V2A11 and V9E1 would further enhance intrabody activity (Herrera *et al*., 2016; Herrera et al., 2015; Vance *et al*., 2013). To test this hypothesis, we engineered a plasmid encoding an N-terminal V2A11 and a C-terminal V9E1, joined via a (G_4_S)_4_ linker of sufficient length to enable simultaneous binding of each V_H_H to RTA (V2A11-V9E1) (**Figure S3, S4**). To confirm the validity of the construct, Vero cells were transfected with the heterodimer-encoding plasmid and resulting cell lysates were examined by Western blot and ELISA. Western blotting of cell lysates with HRP-conjugated, anti-E-tag IgG revealed a polypeptide of ∼ 30 kDa (**Figure S4B**), consistent with the predicted size of the V2A11-V9E1 heterodimer. Moreover, by ELISA, the same Vero cell lysates were reactive with RTA (**Figure S4C)**. Competition assays using epitope specific mAbs or ricin to block one or both of the V2A11 and V9E1 epitopes demonstrated that both V_H_H components of the heterodimer were capable of binding RTA (**Figure S5**).

Vero cells were then transfected with the V2A11-V9E1 encoding plasmid and challenged with increasing amounts of RT. As controls, Vero cells were transfected with V2A11 and V9E1, individually or in combination. The results revealed that Vero cells transfected with the plasmid encoding the V2A11-V9E1 construct were more resistant to RT intoxication compared to vehicle control cells, as evidenced by an RI of 125 (EC_50_ 400 ng/mL vs. 3.2 ng/mL) (**Figure 2**).

**Figure 2.**
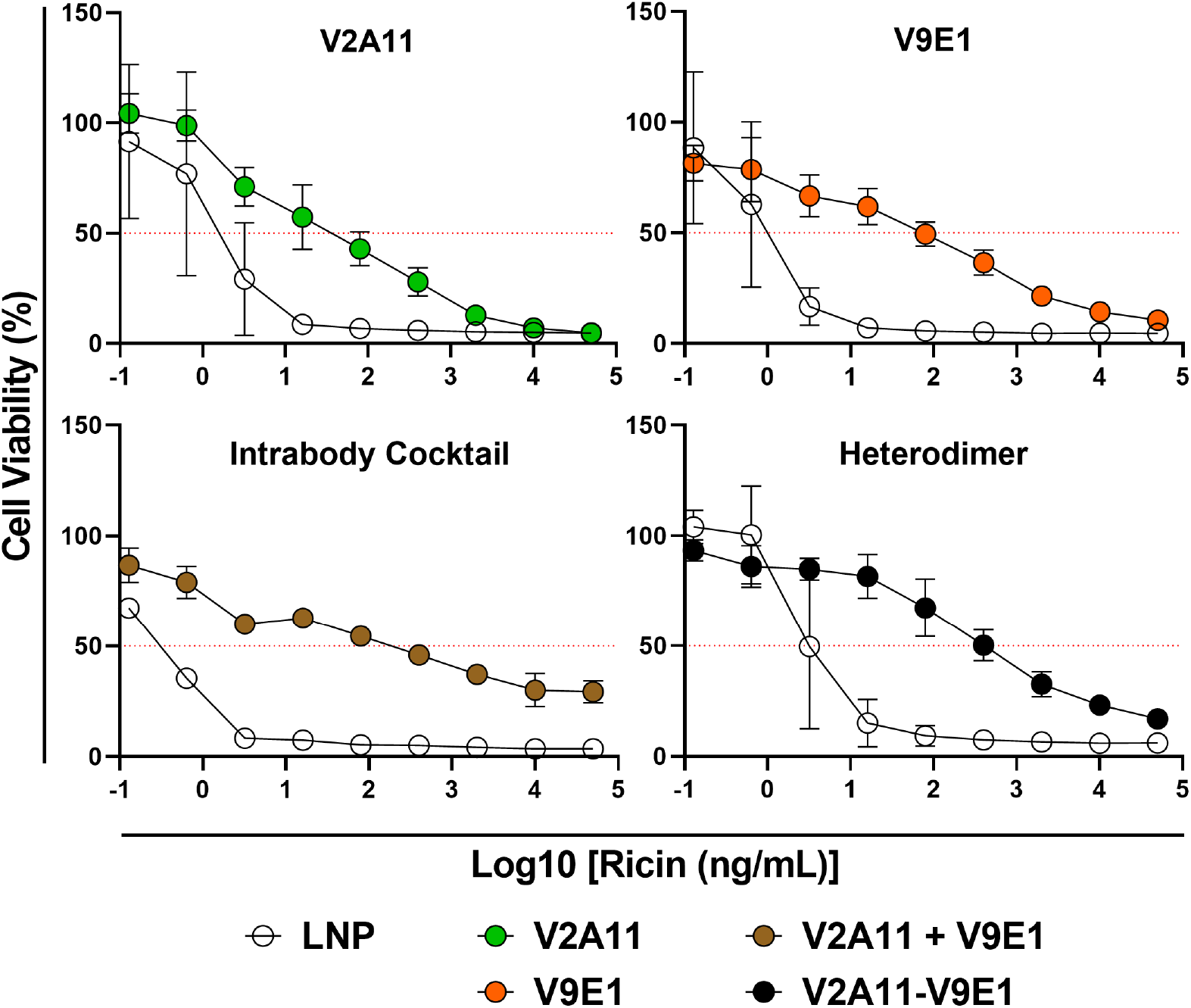
Intracellular expression of V2A11-V9E1 potently protects Vero cells from ricin intoxication. Vero cells were transiently transfected with either V2A11 (green; top left), V9E1 (orange; top right), a cocktail of V2A11 and V9E1 (brown; bottom left), or the V2A11-V9E1 heterodimer (black; bottom right). One day after transfection, cells were treated with a range of ricin concentrations and analyzed for viability two days after ricin treatment. Cell viabilities were normalized to “live” control cells for each transfection condition, not treated with ricin, and compared to cells transfected with vehicle LNP controls (white).

Furthermore, V2A11-V9E1 transfection resulted in RIs of 5 and 11 compared to cells transfected with V9E1 or V2A11, respectively. However, Vero cells transfected with plasmid encoding the covalently linked V2A11-V9E1 construct afforded little benefit as compared to dual transfection with the combination of V2A11 and V9E1 plasmids (**Figure 2**). Given that transient DNA transfection results in only a fraction of the total cells expressing any given V_H_H intrabody, we reasoned that this approach was likely underestimating the full potency of the different intrabodies.

To overcome this issue, we stably transfected Vero cells with DNA constructs encoding GFP-labeled intrabodies and subjected cells to antibiotic selection and fluorescence-based screening. Specifically, Vero cells were transfected with plasmids encoding one of four different V_H_Hs, each with a C-terminal eGFP fusion domain. V9E1 and V2A11-V9E1 were selected based on their potency as intrabodies in transient transfection. V9F6 and ciA-H7, a V_H_H-specific for botulinum neurotoxin, were used as controls. Transfected cells were maintained under Geneticin^®^ (G418) selection. At 6-, 12-, and 15-weeks post-transfection, cells were trypsinized and subjected to fluorescence-activated cell sorting (FACS) to enrich for eGFP+ cells **(Fig S6)**.

For each intrabody, the percentage of fluorescent cells increased at each subsequent FACS purification and only those cells with the highest fluorescence intensity were sorted to select for the highest intrabody expression. Cells transfected with ciA-H7 continued to demonstrate < 1% eGFP positivity and were thus removed from subsequent analyses. Over the next 30 weeks, cells were analyzed by ELISA and ricin cytotoxicity assays to assess intrabody expression and neutralization of the toxin. By ELISA, we determined that levels of intrabody expression was comparable between V9F6-, V9E1-, and V2A11-V9E1-expressing cells (**data not shown**).

At various timepoints, cells were passaged into 96-well plates and treated with serial dilutions of ricin the following day. These cells were analyzed for viability two days after ricin treatment and compared to non-transfected control Vero cells. V9E1-transfected cells were significantly resistant to ricin with an RI of 97 as compared to control cells after 11-weeks and an RI of 192 in experiments performed between 14- and 30-weeks post-transfection (**Figure 3**). At 11 weeks post-transfection, V9F6-expressing cells were moderately resistant to the toxin compared to control cells with an RI of 7, however resistance did not persist beyond this time point.

**Figure 3.**
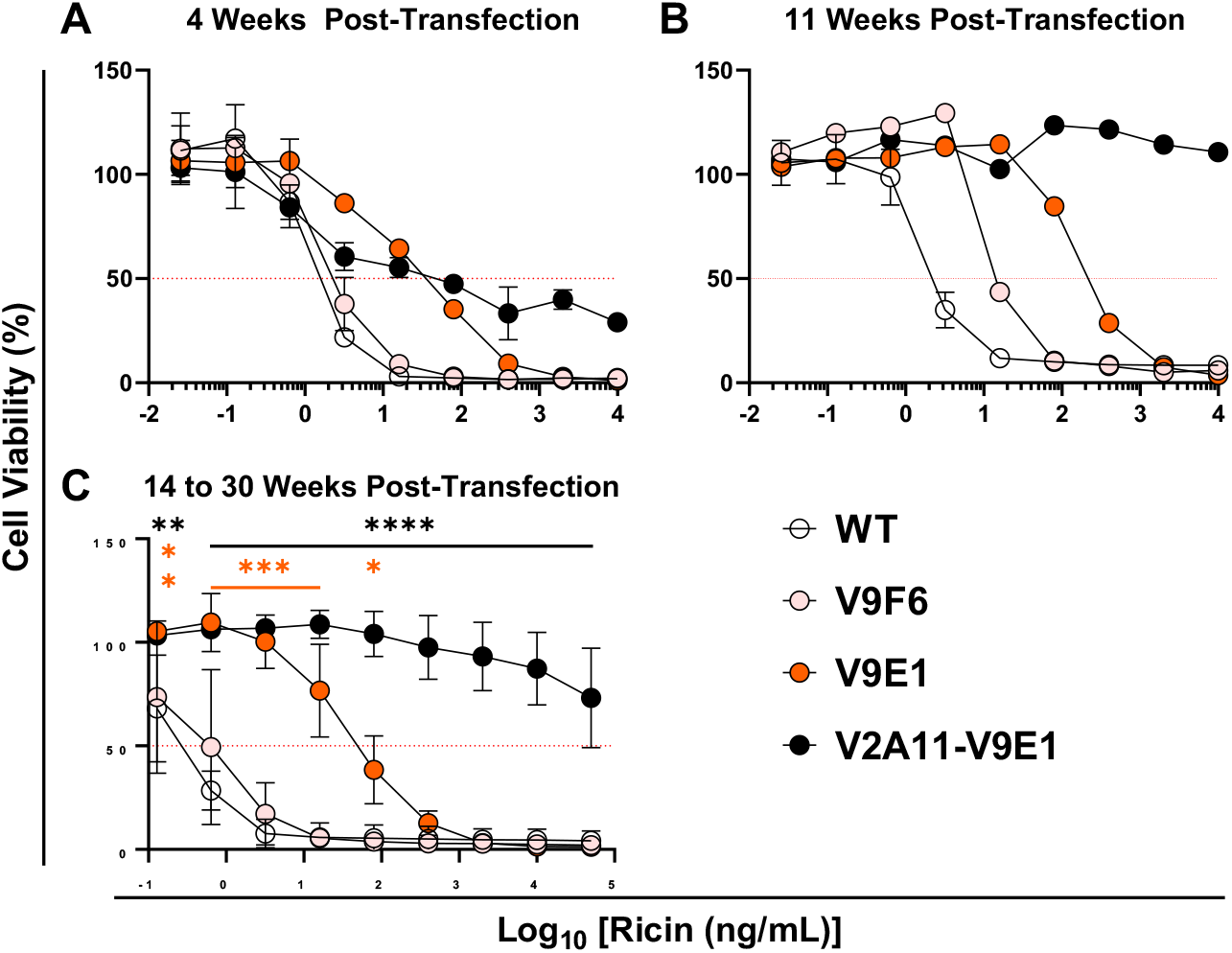
Vero cells stably expressing V2A11-V9E1 are extremely resistant to ricin. Transfected cells were seeded in 96-well plates without selection media and treated with a range of ricin concentrations one day later in triplicate. Cell viabilities were determined two days following ricin treatment, normalized to “live” control cells for each intrabody condition. Un-transfected control Vero cells were grown in DMEM without G418 and tested simultaneously to generate standard ricin dose response curves. Cells were tested in cytotoxicity assays for ricin resistance at A) 4 weeks post-transfection, B) 11 weeks post-transfection, and C) 14-to 30-weeks post-transfection. Throughout the 14-30 weeks post-transfection, cells were continuously subjected to cytotoxicity assays as described above and results averaged. Significance for the 14-to 30-week assays was determined by two-way ANOVA with Sidak post-hoc test at each RTA concentration tested. Black asterisks represent significant differences between V2A11-V9E1-transfected and control Vero cells. Orange asterisks represent significant differences between V9E1 transfected and control cells. **** p<0.0001; *** p<0.001; ** p<0.005; * p<0.05.

Remarkably, cells expressing V2A11-V9E1 were resistant to 10 μg/mL ricin from 11-to 30-weeks post transfection, demonstrating 90-100 % cell viability. At 50 μg/mL of ricin, these cells consistently remained at least 60% viable, representing an RI >250,000, surpassing monomeric V9E1 by a factor of 350 fold (**Figure 3)**. To test whether this resistance was due to the presence of the heterodimer or an off-target effect resulting from the selection process, we removed antibiotic selection from cells expressing the heterodimeric intrabody for extended periods. After 12-weeks, the ricin EC_50_ dropped in these cells from > 50 μg/mL (G418+ media) to 30 ng/mL (G418 removed) (**Figure S7**). This is consistent with cells gradually losing the intrabody-encoding plasmid as they divide in the absence of a selective pressure. Altogether, these results demonstrate that the biparatopic intrabody targeting both the AS and RRS is capable of “ultra-potent” neutralization of ricin when expressed intracellularly, as defined by > 50% cell viability at 50 μg/mL of the toxin.

## Discussion

The CDC’s list of bioterrorism agents represents a diversity of viruses, bacteria and toxins that share the potential to cause morbidity and mortality in otherwise healthy individuals. A report by a panel of biodefense experts considered RT as particularly problematic because of its “….reliability, virulence, and a lack of available countermeasures” (Cieslak *et al*., 2018).

From the standpoint of MCM development, RT is also challenging as it exerts its catalytic activities within the protective confines of the cell cytoplasm where it is off limits to conventional antibody neutralization mechanisms. However, we have shown in previous reports that RTA is susceptible to intracellular neutralization by antibodies targeting the active site (V2A11) or the RRS (V9E1) (Czajka *et al*., 2022; Rudolph *et al*., 2020). In the current study, we have advanced those observations and now demonstrate that the combination of V2A11 and V9E1 as a biparatopic intrabody virtually eliminates RTA’s RIP activity and renders Vero cells impervious to the effects of RT *in vitro*. While animal studies are pending, the prospect of targeting a multimeric (biparatopic) V_H_H into lung tissues as a means to arrest RT intoxication is not farfetched.

Our study adds to the list of examples of engineered biparatopic antibodies that have biological properties (*e*.*g*., toxin neutralization) that exceed what is achieved by either of the variable elements alone, although V2A11-V9E1 may constitute the first reported biparatopic intrabody (Niquille et al., 2024). We postulate that the enhanced potency of V2A11-V9E1 is due to a combination of increased antibody avidity and simultaneous targeting of two sites of vulnerability of RTA. We and others have already described significant increases in antibody avidity associated with bispecific and biparatopic V_H_Hs targeting RT (Glaven et al., 2012; Herrera *et al*., 2016; Vance *et al*., 2013; Vance et al., 2017). Reductions in antibody off rates (k_d_), in particular, correlate with enhanced toxin-neutralizing activity (Vance *et al*., 2017). Perhaps even more consequential is the ability of V2A11-V9E1 to effectively shield the ribosome from RTA by simultaneously occluding both the P-stalk binding pocket (RRS) and the active site (AS). That said we have not provided direct experimental evidence through X-ray crystallography, for example, demonstrating that V2A11 and V9E1 do in fact simultaneously engage their epitopes on a single molecule of RTA. However, the linker length chosen for V2A11-V9E1 is based on structures of other V_H_H-based biparatopic antibodies and is theoretically compatible with dual binding on RTA (Lam et al., 2020).

As a potential therapeutic for inhalational RT intoxication, V2A11-V9E1 would need to be efficiently delivered into targets cells that line the upper and lower airways. While liquid, aerosolized or nebulized delivery of antibodies and nanobodies into the lungs is well established, especially in the wake of the COVID-19 pandemic (Leyva-Grado et al., 2015; Parray et al., 2021; Rong et al., 2020a; Rong et al., 2020b; Streblow et al., 2023), getting them into the intracellular compartment to function as intrabodies is a different matter altogether. However, great strides have been made in the use of mRNA-based platforms for delivery of pathogen and toxin-specific IgG and sdAbs into mucosal tissues, including the lung (Deal et al., 2023; Panova et al., 2023; Tai et al., 2023; Thran et al., 2017). At the same time, methods for optimizing intracellular V_H_H expression have been elucidated (Dingus et al., 2022). Thus, the technological methodologies required to evaluate V9E1 and V2A11, individually or in combination, in a mouse or even non-human primate model are available. The non-human primate model for aerosolized RT intoxication has the advantage of having already been used to evaluate the RTA-specific monoclonal IgG, PB10, in a pre- and post-exposure model (Roy *et al*., 2019; Roy *et al*., 2020). PB10 targets an immunodominant epitope near RTA’s active site, but, by all accounts, neutralizes RT by engaging with the toxin extracellularly and interfering with uptake and retrograde transport (Yermakova et al., 2016). Thus, when administered intravenously to non-human primates, we proposed that PB10 accesses the airways by transudation and intercepts the toxin prior to entry into alveolar cells. Accordingly, PB10 was only effective at rescuing intoxicated animals within 4 h of toxin exposure (Roy *et al*., 2019). We speculate that the addition of the V9E1-V2A11 biparatopic intrabody would extend that therapeutic window and reduce mortality and morbidity. In the case of botulinum neurotoxin (BoNT), intracellular targeting of VHHs were able to reverse the toxin’s lethal effects (Miyashita et al., 2021; Wu et al., 2021).

## Materials and Methods

### Construction of mammalian VHH expression plasmids

gBlocks™ (Integrated DNA Technologies, Coralville, IA) encoding V_H_H were ligated into the pcDNA3.1 mammalian expression vector (Addgene, Watertown, MA), in frame with a carboxy terminal E-tag or eGFP-tag for transient or stable transfection, respectively(Czajka and Mantis, 2022; Tremblay et al., 2010).

### Vero Cell Transfection

Vero cells (ATCC, Manassas, VA) were maintained in Dulbecco’s minimal essential medium (DMEM) with fetal bovine serum (10% v/v) and penicillin/streptomycin (referred to henceforth simply as DMEM) at 37°C (5% CO_2_). For transient transfection experiments, cells were seeded one day prior to transfection in 6-or 96-well plates at a density of 10^5^ cells/mL, as recommended by the Lipofectamine™ 3000 transfection protocol (Life Technologies, Carlsbad, CA). Cells were transfected with 250 μL/well (6-well plate) or 10 μL/well (96-well plate) of transfection mixtures containing intrabody DNA plasmids according to the recommended Lipofectamine™ 3000 transfection protocol.

### Extraction of cell lysate from transfected cells

Cells were washed 48 h post-transfection twice in ice-cold PBS. RIPA Lysis Buffer (50 mM Tris-HCl pH7.5, 150 mM NaCl, 1% NP-40, 0.1% SDS, and 1% Na-deoxycholate; 150 μL) was added and cells were incubated for 5 minutes on ice with occasional gentle rocking to ensure well coverage. Cells were transferred to chilled 2 mL screw cap tubes with 1 mm glass beads by scraping and pipetting followed by homogenization at 5 m/s for 5 s. Cell lysates were replaced on ice for 30 min with one more homogenization step, followed by centrifugation at 15,700 × g for 10 min at 4°C.

### Cell lysate ELISA

96-well ELISA plates were coated overnight at 4°C with 1 μg/mL PH12 in PBS. Plates were washed and blocked for 2 h at room temperature. Following block, RTA (1 μg/mL in PBS) was applied to ELISA plates for 1 h. Transfected cell lysate was serially diluted in duplicate and added to plates for 1 h. Plates were washed and anti-E-HRP antibody(Bethyl Laboratories; 1:10,000) was applied for 1 h. For e-GFP-Fusion intrabodies, V_H_Hs in lysate were detected via C-terminal eGFP domains with polyclonal rabbit anti-eGFP antibody (ThermoFisher; 1:5,000 in Wash Buffer) followed by HRP-conjugated goat anti-rabbit secondary antibody (SouthernBiotech; 1:2,000 in Wash Buffer). Plates were washed and 100 μL TMB was added for 5-10 min followed by 100 µL Stop Solution. ELISA plates were analyzed using a SpectraMax iD3 spectrophotometer equipped with Softmax Pro 7 software at OD_450_. Purified V_H_H protein with a C-terminal E-tag was used as a positive binding control for each transfection. Plates were washed following each step in PBS-Tween (0.1%). Cell lysates and secondary antibody were diluted in block buffer (PBS-Tween supplemented with goat serum (10%).

For lysate from Vero cells stably expressing the V2A11-V9E1 heterodimer, plates were coated overnight at 4°C with either 1 µg/mL PH12 or IB2. Plates were blocked as described above. RTA (1 µg/mL), RTA+V9B2 (10 µg/mL), or ricin (1 µg/mL) was added for 1 h at room temperature. Plates were washed and pooled cell lysate was serially diluted and added. Intrabody binding was detected via C-terminal eGFP as described above.

### Western Blot

Cell lysates were diluted into loading buffer, boiled, and separated by electrophoresis on precast 4%–15% polyacrylamide gradient gels (Bio-Rad Laboratories, Herculues, CA). Separated proteins were transferred to nitrocellulose membranes and then blocked overnight at 4°C in Western blotting block buffer (TBS-Tween (0.5 %) with 5% BSA). HRP-conjugated anti-E-tag antibody (1:10,000) was added for 1 h at room temperature with gentle rocking. Membranes were washed (TBS-T) three times for 10 min at room temperature with gentle rocking. Samples were detected with the Pierce ECL Plus Detection Kit (ThermoFisher) and using the iBright Imager set to “Universal Mode” to visualize AlexaFluor647 and ECL channels for fluorescent ladder and samples, respectively.

### Ricin cytotoxicity assay

One day after transfection in 96-well plates, medium was aspirated from cells, replaced with 100 μL ricin toxin serially diluted in DMEM, and incubated at 37°C for 2 h. Ricin was then aspirated from cells and replaced with 100 μL DMEM. Cells were incubated for 48 h at 37°C. Viability was determined using Cell Titer-Glo® (Promega) and a SpectraMax L or SpectraMax iD3 Microplate Reader to read luminescence and measured as a percentage of live control cells (transfected, but not treated with ricin).

For cytotoxicity assays with stably transfected cells, cells were split and seeded in 96-well plates in cell culture media without G418 at a density of 10^5^ cells/mL. One day after seeding, cells were treated with serial dilutions of ricin as described previously for transient-intrabody ricin cytotoxicity assays. Control Vero cells not transfected or grown in G418 supplemented media were seeded at the same density and treated with ricin to generate standard dose-response curves. For experiments in which G418 selection was removed, these cells were passaged in normal cell culture medium (DMEM + 10% FBS + 1X P/S).

### Stable cell line generation

Vero cells were seeded in a 6-well plate at 10^5^ cells/mL and transfected with pcDNA3 plasmids encoding V_H_Hs with C-terminal eGFP fusion domains. Two days post-transfection, cells were passaged and diluted 1:125 in cell culture media containing 600 ug/mL G418 (“Selection Media”). Cells were kept under selective pressure for three weeks with Selection Media replaced every 2-3 days until mock transfected control cells were dead and colonies formed for each intrabody condition.

Cells were passaged in cell culture media containing 400 ug/mL G418 (“Maintenance Media”) with media refreshed every 2-3 days. After 2 weeks, eGFP+ cells were sorted into 48-well tissue culture plates using the BD FACSAria™ II Cell Sorter (BD Biosciences, Franklin Lakes, NJ) and cultured in Maintenance Media. Media was collected and analyzed for the presence of V_H_Hs via ELISA, as described below. Wells with the highest RTA-binding (OD450) for each intrabody cell line were selected and cells were grown in T75 flasks and re-sorted based on highest median fluorescence intensity into 6-well tissue culture plates. Cells were passaged and analyzed for intracellular V_H_H expression via ELISA and those with the highest RTA-binding for each intrabody condition were selected for the final round of sorting.

### Statistical analysis

Statistical analyses were performed using GraphPad Prism 8.2 or 9.1 software for Windows (San Diego, California). For intrabody cytotoxicity results, a two-way ANOVA was used with the Sidak post hoc test to compare transfected and LNP-vehicle control cell viabilities at each ricin concentration. To predict intrabody-based Vero cell protection, a multiple linear regression least squares analysis was performed with cytotoxicity area under the curve as the dependent outcome and intrabody ELISA and V9B2 competition areas under the curve as independent variables.

### V2A11-V9E11 heterodimer modeling

Structural alignments of V2A11-RTA (PDB ID 6OBC) and V9E1-RTA (PDB ID 7TGI) complexes were generated using the PyMOL Molecular Graphics System, Version 2.0. The distance between V2A11 and V9E1 was approximated between the C-terminal residue Ser122 of V2A11 and the most N-terminal residue of the V9E1 structure, Leu4.

## Author contributions

TFC: Conceptualization, Methodology, Investigation, Formal Analysis, Writing – Original Draft, Writing – Review and Editing, Visualization. DJV: Conceptualization, Writing – Review and Editing, Supervision. RS: Methodology, Investigation. NJM: Conceptualization, Writing – Review and Editing, Supervision, Project Administration, Funding Acquisition.

## Data availability

All data associated with this study are included in the manuscript.

## Acknowledgements

We gratefully acknowledge Dr. Charles Shoemaker (Tufts University) for his instrumental insight into the design and expression of V_H_Hs. We thank Elizabeth Cavosie (Wadsworth Center) for administrative assistance.

## References Cited

Anderson, G.P., Liu, J.L., Hale, M.L., Bernstein, R.D., Moore, M., Swain, M.D., and Goldman, E.R. (2008). Development of antiricin single domain antibodies toward detection and therapeutic reagents. Anal Chem 80, 9604–9611. 10.1021/ac8019398.

Chiou, J.C., Li, X.P., Remacha, M., Ballesta, J.P., and Tumer, N.E. (2008). The ribosomal stalk is required for ribosome binding, depurination of the rRNA and cytotoxicity of ricin A chain in Saccharomyces cerevisiae. Mol Microbiol 70, 1441–1452. 10.1111/j.1365-2958.2008.06492.x.

Cieslak, T.J., Kortepeter, M.G., Wojtyk, R.J., Jansen, H.J., Reyes, R.A., Smith, J.O., and And the, N.B.M.A.P. (2018). Beyond the Dirty Dozen: A Proposed Methodology for Assessing Future Bioweapon Threats. Mil Med 183, e59–e65. 10.1093/milmed/usx004.

Czajka, T.F., and Mantis, N.J. (2022). Single-Domain Antibodies for Intracellular Toxin Neutralization. Methods Mol Biol 2446, 469–487. 10.1007/978-1-0716-2075-5_24.

Czajka, T.F., Vance, D.J., Davis, S., Rudolph, M.J., and Mantis, N.J. (2022). Single-domain antibodies neutralize ricin toxin intracellularly by blocking access to ribosomal P-stalk proteins. J Biol Chem 298, 101742. 10.1016/j.jbc.2022.101742.

Deal, C.E., Richards, A.F., Yeung, T., Maron, M.J., Wang, Z., Lai, Y.T., Fritz, B.R., Himansu, S., Narayanan, E., Liu, D., et al. (2023). An mRNA-based platform for the delivery of pathogen-specific IgA into mucosal secretions. Cell Rep Med 4, 101253. 10.1016/j.xcrm.2023.101253.

Dingus, J.G., Tang, J.C.Y., Amamoto, R., Wallick, G.K., and Cepko, C.L. (2022). A general approach for stabilizing nanobodies for intracellular expression. Elife 11. 10.7554/eLife.68253.

Endo, Y., and Tsurugi, K. (1987). RNA N-glycosidase activity of ricin A-chain. Mechanism of action of the toxic lectin ricin on eukaryotic ribosomes. J Biol Chem 262, 8128–8130.

Endo, Y., and Tsurugi, K. (1988). The RNA N-glycosidase activity of ricin A-chain. The characteristics of the enzymatic activity of ricin A-chain with ribosomes and with rRNA. J Biol Chem 263, 8735–8739.

Gal, Y., Mazor, O., Falach, R., Sapoznikov, A., Kronman, C., and Sabo, T. (2017). Treatments for Pulmonary Ricin Intoxication: Current Aspects and Future Prospects. Toxins (Basel) 9, 311. 10.3390/toxins9100311.

Glaven, R.H., Anderson, G.P., Zabetakis, D., Liu, J.L., Long, N.C., and Goldman, E.R. (2012). Linking Single Domain Antibodies that Recognize Different Epitopes on the Same Target. Biosensors (Basel) 2, 43–56. 10.3390/bios2010043.

Herrera, C., Klokk, T.I., Cole, R., Sandvig, K., and Mantis, N.J. (2016). A Bispecific Antibody Promotes Aggregation of Ricin Toxin on Cell Surfaces and Alters Dynamics of Toxin Internalization and Trafficking. PLoS One 11, e0156893. 10.1371/journal.pone.0156893.

Herrera, C., Tremblay, J.M., Shoemaker, C.B., and Mantis, N.J. (2015). Mechanisms of Ricin Toxin Neutralization Revealed through Engineered Homodimeric and Heterodimeric Camelid Antibodies. J Biol Chem 290, 27880–27889. 10.1074/jbc.M115.658070.

Lam, K.H., Tremblay, J.M., Vazquez-Cintron, E., Perry, K., Ondeck, C., Webb, R.P., McNutt, P.M., Shoemaker, C.B., and Jin, R. (2020). Structural Insights into Rational Design of Single-Domain Antibody-Based Antitoxins against Botulinum Neurotoxins. Cell Rep 30, 2526–2539 e2526. 10.1016/j.celrep.2020.01.107.

Laursen, N.S., Friesen, R.H.E., Zhu, X., Jongeneelen, M., Blokland, S., Vermond, J., van Eijgen, A., Tang, C., van Diepen, H., Obmolova, G., et al. (2018). Universal protection against influenza infection by a multidomain antibody to influenza hemagglutinin. Science 362, 598–602. 10.1126/science.aaq0620.

Leyva-Grado, V.H., Tan, G.S., Leon, P.E., Yondola, M., and Palese, P. (2015). Direct administration in the respiratory tract improves efficacy of broadly neutralizing anti-influenza virus monoclonal antibodies. Antimicrob Agents Chemother 59, 4162–4172. 10.1128/AAC.00290-15.

Li, X.P., Chiou, J.C., Remacha, M., Ballesta, J.P., and Tumer, N.E. (2009). A two-step binding model proposed for the electrostatic interactions of ricin a chain with ribosomes. Biochemistry 48, 3853–3863. 10.1021/bi802371h.

Li, X.P., Kahn, J.N., and Tumer, N.E. (2018). Peptide Mimics of the Ribosomal P Stalk Inhibit the Activity of Ricin A Chain by Preventing Ribosome Binding. Toxins (Basel) 10. 10.3390/toxins10090371.

May, K.L., Li, X.P., Martinez-Azorin, F., Ballesta, J.P., Grela, P., Tchorzewski, M., and Tumer, N.E. (2012). The P1/P2 proteins of the human ribosomal stalk are required for ribosome binding and depurination by ricin in human cells. FEBS J 279, 3925–3936. 10.1111/j.1742-4658.2012.08752.x.

Miyashita, S.I., Zhang, J., Zhang, S., Shoemaker, C.B., and Dong, M. (2021). Delivery of single-domain antibodies into neurons using a chimeric toxin-based platform is therapeutic in mouse models of botulism. Sci Transl Med 13. 10.1126/scitranslmed.aaz4197.

Mlsna, D., Monzingo, A.F., Katzin, B.J., Ernst, S., and Robertus, J.D. (1993). Structure of recombinant ricin A chain at 2.3 A. Protein Sci 2, 429–435.

Montfort, W., Villafranca, J.E., Monzingo, A.F., Ernst, S.R., Katzin, B., Rutenber, E., Xuong, N.H., Hamlin, R., and Robertus, J.D. (1987). The three-dimensional structure of ricin at 2.8 A. Journal of Biological Chemistry. 262, 5398–5403.

Muyldermans, S. (2021). A guide to: generation and design of nanobodies. FEBS J 288, 2084–2102. 10.1111/febs.15515.

Niquille, D.L., Fitzgerald, K.M., and Gera, N. (2024). Biparatopic antibodies: therapeutic applications and prospects. MAbs 16, 2310890. 10.1080/19420862.2024.2310890.

Olsnes, S., Refsnes, K., and Pihl, A. (1974). Mechanism of action of the toxic lectins abrin and ricin. Nature 249, 627–631. 10.1038/249627a0.

Panova, E.A., Kleymenov, D.A., Shcheblyakov, D.V., Bykonia, E.N., Mazunina, E.P., Dzharullaeva, A.S., Zolotar, A.N., Derkaev, A.A., Esmagambetov, I.B., Sorokin, II, et al. (2023). Single-domain antibody delivery using an mRNA platform protects against lethal doses of botulinum neurotoxin A. Front Immunol 14, 1098302. 10.3389/fimmu.2023.1098302.

Parray, H.A., Shukla, S., Perween, R., Khatri, R., Shrivastava, T., Singh, V., Murugavelu, P., Ahmed, S., Samal, S., Sharma, C., et al. (2021). Inhalation monoclonal antibody therapy: a new way to treat and manage respiratory infections. Appl Microbiol Biotechnol 105, 6315–6332. 10.1007/s00253-021-11488-4.

Pincus, S.H., Bhaskaran, M., Brey, R.N., 3rd, Didier, P.J., Doyle-Meyers, L.A., and Roy, C.J. (2015). Clinical and Pathological Findings Associated with Aerosol Exposure of Macaques to Ricin Toxin. Toxins (Basel) 7, 2121–2133. 10.3390/toxins7062121.

Poon, A.Y., Vance, D.J., Rong, Y., Ehrbar, D., and Mantis, N.J. (2017). A Supercluster of Neutralizing Epitopes at the Interface of Ricin’s Enzymatic (RTA) and Binding (RTB) Subunits. Toxins (Basel) 9. 10.3390/toxins9120378.

Raybould, M.I.J., Kovaltsuk, A., Marks, C., and Deane, C.M. (2021). CoV-AbDab: the coronavirus antibody database. Bioinformatics 37, 734–735. 10.1093/bioinformatics/btaa739.

Rong, Y., Pauly, M., Guthals, A., Pham, H., Ehrbar, D., Zeitlin, L., and Mantis, N.J. (2020a). A Humanized Monoclonal Antibody Cocktail to Prevent Pulmonary Ricin Intoxication. Toxins (Basel) 12, 215. 10.3390/toxins12040215.

Rong, Y., Torres-Velez, F.J., Ehrbar, D., Doering, J., Song, R., and Mantis, N.J. (2020b). An intranasally administered monoclonal antibody cocktail abrogates ricin toxin-induced pulmonary tissue damage and inflammation. Hum Vaccin Immunother 16, 793–807. 10.1080/21645515.2019.1664243.

Roy, C.J., Ehrbar, D.J., Bohorova, N., Bohorov, O., Kim, D., Pauly, M., Whaley, K., Rong, Y., Torres-Velez, F.J., Vitetta, E.S., et al. (2019). Rescue of rhesus macaques from the lethality of aerosolized ricin toxin. JCI Insight 4. 10.1172/jci.insight.124771.

Roy, C.J., Van Slyke, G., Ehrbar, D., Bornholdt, Z.A., Brennan, M.B., Campbell, L., Chen, M., Kim, D., Mlakar, N., Whaley, K.J., et al. (2020). Passive immunization with an extended half-life monoclonal antibody protects Rhesus macaques against aerosolized ricin toxin. NPJ Vaccines 5, 13. 10.1038/s41541-020-0162-0.

Rudolph, M.J., Czajka, T.F., Davis, S.A., Thi Nguyen, C.M., Li, X.P., Tumer, N.E., Vance, D.J., and Mantis, N.J. (2020). Intracellular Neutralization of Ricin Toxin by Single-domain Antibodies Targeting the Active Site. J Mol Biol 432, 1109–1125. 10.1016/j.jmb.2020.01.006.

Rudolph, M.J., Poon, A.Y., Kavaliauskiene, S., Myrann, A.G., Reynolds-Peterson, C., Davis, S.A., Sandvig, K., Vance, D.J., and Mantis, N.J. (2021). Structural Analysis of Toxin-Neutralizing, Single-Domain Antibodies that Bridge Ricin’s A-B Subunit Interface. J Mol Biol 433, 167086. 10.1016/j.jmb.2021.167086.

Rutenber, E., Katzin, B.J., Ernst, S., Collins, E.J., Mlsna, D., Ready, M.P., and Robertus, J.D. (1991). Crystallographic refinement of ricin to 2.5 A. Proteins 10, 240–250. 10.1002/prot.340100308.

Snieckute, G., Genzor, A.V., Vind, A.C., Ryder, L., Stoneley, M., Chamois, S., Dreos, R., Nordgaard, C., Sass, F., Blasius, M., et al. (2022). Ribosome stalling is a signal for metabolic regulation by the ribotoxic stress response. Cell Metab 34, 2036–2046 e2038. 10.1016/j.cmet.2022.10.011.

Sowa-Rogozinska, N., Sominka, H., Nowakowska-Golacka, J., Sandvig, K., and Slominska-Wojewodzka, M. (2019). Intracellular Transport and Cytotoxicity of the Protein Toxin Ricin. Toxins (Basel) 11. 10.3390/toxins11060350.

Streblow, D.N., Hirsch, A.J., Stanton, J.J., Lewis, A.D., Colgin, L., Hessell, A.J., Kreklywich, C.N., Smith, J.L., Sutton, W.F., Chauvin, D., et al. (2023). Aerosol delivery of SARS-CoV-2 human monoclonal antibodies in macaques limits viral replication and lung pathology. Nat Commun 14, 7062. 10.1038/s41467-023-42440-x.

Tai, W., Yang, K., Liu, Y., Li, R., Feng, S., Chai, B., Zhuang, X., Qi, S., Shi, H., Liu, Z., et al. (2023). A lung-selective delivery of mRNA encoding broadly neutralizing antibody against SARS-CoV-2 infection. Nat Commun 14, 8042. 10.1038/s41467-023-43798-8.

Thran, M., Mukherjee, J., Ponisch, M., Fiedler, K., Thess, A., Mui, B.L., Hope, M.J., Tam, Y.K., Horscroft, N., Heidenreich, R., et al. (2017). mRNA mediates passive vaccination against infectious agents, toxins, and tumors. EMBO Mol Med 9, 1434–1447. 10.15252/emmm.201707678.

Tremblay, J.M., Kuo, C.L., Abeijon, C., Sepulveda, J., Oyler, G., Hu, X., Jin, M.M., and Shoemaker, C.B. (2010). Camelid single domain antibodies (VHHs) as neuronal cell intrabody binding agents and inhibitors of Clostridium botulinum neurotoxin (BoNT) proteases. Toxicon 56, 990–998. 10.1016/j.toxicon.2010.07.003.

Vance, D.J., Poon, A.Y., and Mantis, N.J. (2020). Sites of vulnerability on ricin B chain revealed through epitope mapping of toxin-neutralizing monoclonal antibodies. PLoS One 15, e0236538. 10.1371/journal.pone.0236538.

Vance, D.J., Tremblay, J.M., Mantis, N.J., and Shoemaker, C.B. (2013). Stepwise engineering of heterodimeric single domain camelid VHH antibodies that passively protect mice from ricin toxin. J Biol Chem 288, 36538–36547. 10.1074/jbc.M113.519207.

Vance, D.J., Tremblay, J.M., Rong, Y., Angalakurthi, S.K., Volkin, D.B., Middaugh, C.R., Weis, D.D., Shoemaker, C.B., and Mantis, N.J. (2017). High-Resolution Epitope Positioning of a Large Collection of Neutralizing and Nonneutralizing Single-Domain Antibodies on the Enzymatic and Binding Subunits of Ricin Toxin. Clin Vaccine Immunol 24, e00236–00217. 10.1128/CVI.00236-17.

Wu, X., Cheng, L., Fu, M., Huang, B., Zhu, L., Xu, S., Shi, H., Zhang, D., Yuan, H., Nawaz, W., et al. (2021). A potent bispecific nanobody protects hACE2 mice against SARS-CoV-2 infection via intranasal administration. Cell Rep 37, 109869. 10.1016/j.celrep.2021.109869.

Yermakova, A., Klokk, T.I., O’Hara, J.M., Cole, R., Sandvig, K., and Mantis, N.J. (2016). Neutralizing Monoclonal Antibodies against Disparate Epitopes on Ricin Toxin’s Enzymatic Subunit Interfere with Intracellular Toxin Transport. Sci Rep 6, 22721. 10.1038/srep22721.

